# Agnostic material classification using differential de Bruijn graphs of DNA imprints

**DOI:** 10.64898/2026.06.23.733838

**Authors:** Rachael M. Cox, Zoya T. Ansari, Charlie D. Johnson, Edward M. Marcotte, Andrew D Ellington, Sanchita Bhadra

## Abstract

The wide variety of physical and chemical properties in materials makes the study of unknown substances challenging. We have previously proposed a theoretical framework for agnostic material characterization based on using nucleic acid ‘imprints’ of the materials and then analyzing material-specific patterns of derived sequences. Here we demonstrate an experimental and computational pipeline that can agnostically identify and distinguish varied materials based on DNA *k*-mer imprints and validate the ability of these imprints to distinguish closely related materials. This work lays the foundation for expansion of purely agnostic sensing technologies for the unbiased characterization and categorization of a much wider variety of biotic and abiotic materials.

## Introduction

Non-Terran life may be composed of molecules that are largely distinct from Terran biology, and thus there is a need for the development of methods for ‘agnostic biosignature’ discovery and detection.^1,2^ Such methods should not only enable ‘search-for-life’ endeavors in future space missions but can also potentially support efforts in molecular diagnostics and materials analyses. The question of how to achieve agnostic biosensing has previously been considered, and possible options include both planet-scale remote sensing or biochemical material analyses that do not rely on detection / presence of known life-associated characteristics or polymers.^3^ Several remote sensing biosignatures have been explored including formation of higher than geologically expected levels of methane hazes in anoxic environments, presence of volatile molecules and seasonality in gas abundance, and existence of unusual global disequilibria such as simultaneous presence of oxygen and methane gases.^3^ Differences in the statistical complexity of temporal variability of planetary light emission or reflection have also been applied as potential agnostic measures of exoplanetary complexity, which in turn could be considered to be indicative of the presence or absence of a biosphere.^4^

While many biochemical approaches to life detection have focused on measuring signatures of known biology or biological activity, such as biomolecules, isotopic patterns, chirality, and charge distribution, a few agnostic approaches that correlate molecular complexity to biotic compounds have also been described. Pyrolysis of terrestrial and non-terrestrial carbonaceous compounds followed by gas chromatography–mass spectrometry (pyro-GC-MS) has been used to develop a machine learning-based classification model that demonstrated 90% accuracy in differentiating known biotic versus abiotic materials.^5^ Similarly, Cronin and co-workers have developed a so-called molecular assembly (MA) index, an integer that represents the shortest synthetic pathway of a molecule from its simpler constituents, which was measured for millions of molecules using mass spectrometry and shown to be an agnostic indicator of molecular complexity and biogenesis.^6^

We have previously proposed that ‘imprinting’ nucleic acids on analytes or materials, followed by DNA sequencing, might yield a complex signature that could be used to characterize living systems that differ from our own.^7^ DNA libraries have previously been sieved for sequences and structures that adhere to a wide variety of materials including natural and chemical polymers, small molecules, minerals and metals.^7,8^ Multiple rounds of selection can yield DNA structures similar to protein antibodies, and such aptamers bind with considerable affinity and specificity even in complex milieus. Thousands of aptamers have been described to bind specific target molecules including proteins, small molecules, and carbohydrates.^9^

However, aptamers against a known set of analytes would not necessarily prove useful in extraterrestrial contexts, especially if living systems were otherwise quite different from our own. That said, the chemistry seen for initial interactions, the fact that nucleic acids can bind and distinguish a wide variety of analytes, can potentially be exploited for agnostic biosensing. We thus propose to briefly sieve large libraries for simpler sequence imprints that might prove to be an agnostic ‘reflection’ of chemistry or structure of analytes and materials. In a similar vein, studies have found that a single round of selection, followed by high-throughput sequencing and *k*-mer analysis, can identify aptamers against targets.^10^ Likewise, mixtures of previously selected aptamers could be used for distinguishing cell lines based on differential patterns and counts of cell-associated sequences.^11^ In fact, even without pre-supposing or knowing the principal targets for aptamer selection, nucleic acid libraries can be enriched for a population of binders that recognizes complex patterns such as different cell surfaces allowing identification of cellular subpopulations in complex mixtures,^12^ tumor-associated biomarkers enabling distinction of treatment-responsive from unresponsive breast cancer,^13^ and changes in exosome composition indicating presence or absence of cancer.^14^

To further advance the hypothesis that we can in general ‘imprint’ chemistry or structure of analytes and materials onto nucleic acids, and recognize sequence motifs and fingerprints of the imprinted molecules, we have now developed a computational pipeline that reconstructs known DNA imprints of oligonucleotide analytes from complex datasets, and have further applied this pipeline to recognize otherwise unknown biotic and abiotic materials such as streptavidin, borosilicate, and chitin. This work lays the foundation for future unbiased categorization of a wide variety of compounds and surfaces, whether biotic or abiotic.

## Results and Discussion

We initially wanted to generate imprints of known nucleic acid sequences, as this would allow us to best characterize our analytical pipeline prior to applying it to unknown materials. To this end, a 90 nucleotide long single-stranded DNA library that contained 15 consecutive randomized nucleotides (N_15_ ssDNA), flanked on the 5’ and 3’ ends by constant sequences of 32 and 43 nucleotides, respectively, was synthesized. Some two picomoles (1.2 x 10^12^ molecules) of this chemically synthesized, unamplified library was applied to four discrete biotinylated hemiduplex DNA substrates, termed Oligo 1, Oligo 2, Oligo 3, or Oligo 4, immobilized separately on the surface of streptavidin-magnetic beads (**Figure 1**).

**Figure 1.**
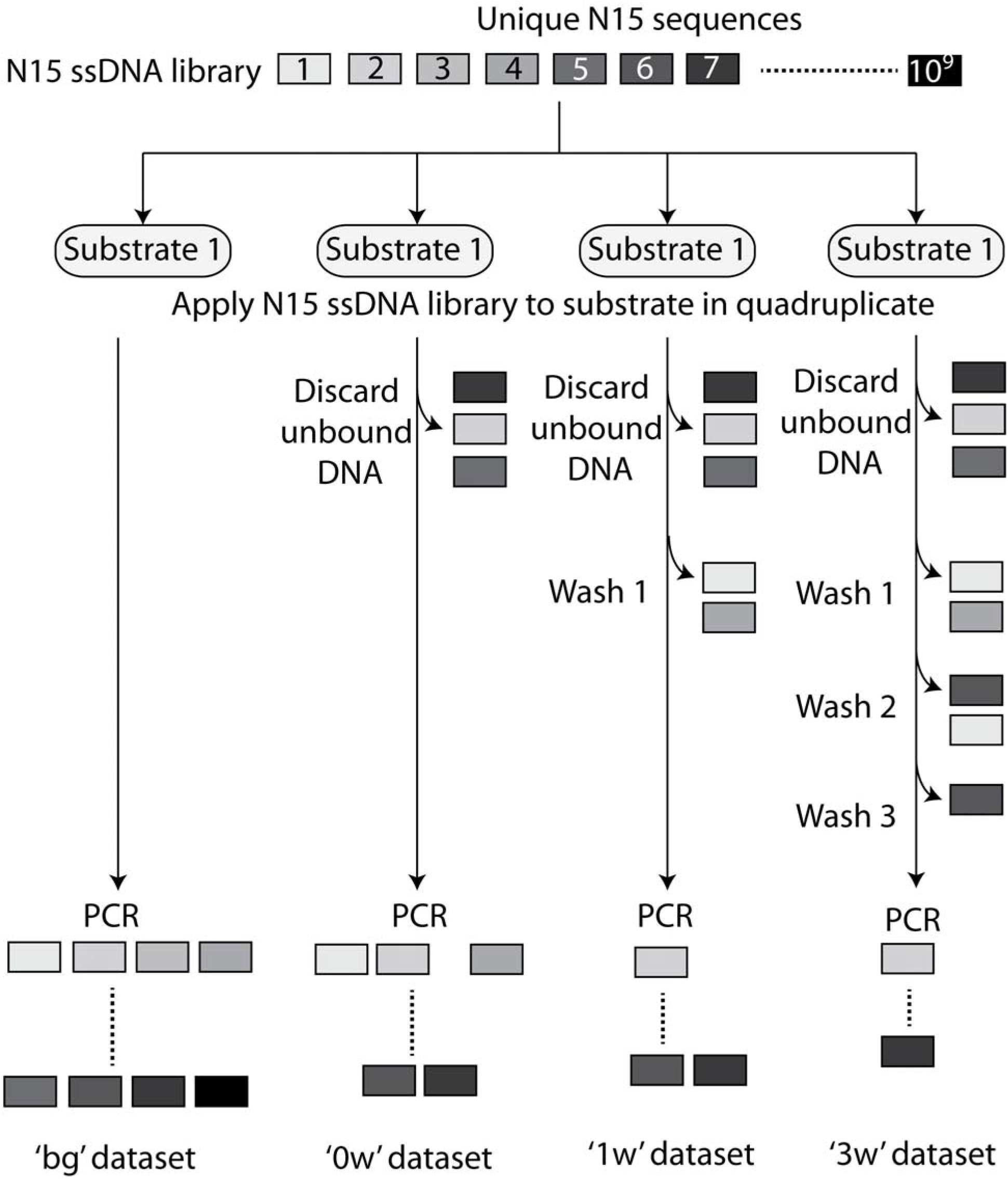
Schematic depicting experimental set up for capturing the sequence imprint of differential substrate binding by a N_15_ single stranded DNA library.

Each oligonucleotide substrate type was incubated with the N_15_ ssDNA library in quadruplicate. The first substrate sample, along with its entire allotment of the N_15_ ssDNA library (termed ‘bg’), was added directly to PCR reactions without further treatment. In the second substrate sample, termed ‘0w’, the supernatant was discarded, and substrate-bound library members were then amplified by adding the substrate to a PCR amplification reaction. In the third (‘1w’) and fourth (‘3w’) substrate samples, supernatant was removed and the samples were washed either once (‘1w’) or thrice (‘3w’) prior to adding the substrates to their respective PCR reactions. These wash conditions had been chosen based on retention of fluorophore-labeled single-stranded DNAs that were fully-, partially-, or non-complementary to bead-bound oligonucleotide targets under different wash conditions (**Supplementary Figure 1**).

All amplicons from PCR reactions were sequenced by massively parallel, paired-end sequencing-by-synthesis using the Illumina platform. The resultant NextGen sequencing datasets were analyzed to determine whether there were substrate-specific patterns in the enrichment of N_15_ sequence motifs. It should be noted that ‘imprinting’ the library on the target molecules differed from a typical aptamer selection, in that there was no intent to drive the pool towards only a relatively few, very high affinity sequences. Thus, the sequence data derived is expected to still be highly diverse, and a key question that these studies address is whether and how sequence imprints can still be derived from such highly noisy, lightly selected populations.

Sequenced populations were analyzed in terms of short subsegments (*k*-mers) within the N_15_ region that ranged from 5 to 15 nt in length. As a first step, the paired end reads for each sequencing dataset were converted to directed edge-centric de Bruijn graphs^15^ for *k* = [5:15] with a bespoke computational pipeline (described in **Figure 2** and the **Methods** section). Briefly, each sequence read was broken into linear *k*-mers, starting at the 5’-end with successive *k*-mers overlapping their predecessor by *k*-1 nucleotide. In this manner 11 .csv files containing *k*-mers of increasing length ranging from 5 to 15 nucleotides were generated from each sequence dataset. Those *k*-mers that were clearly identical to sequences in the static 5’ and 3’ region flanks of the N_15_ library were subtracted (**Figure 2**). The remaining N_15_ region *k*-mer sequences that differed from one another at ≥1 nucleotide position(s) were classed as unique and enumerated for each selection condition to evaluate the proportion of expected sequence space (4*^k^*) at each *k*-mer length that was represented by the observed number of unique *k*-mers (**Figure 3A**).

**Figure 2.**
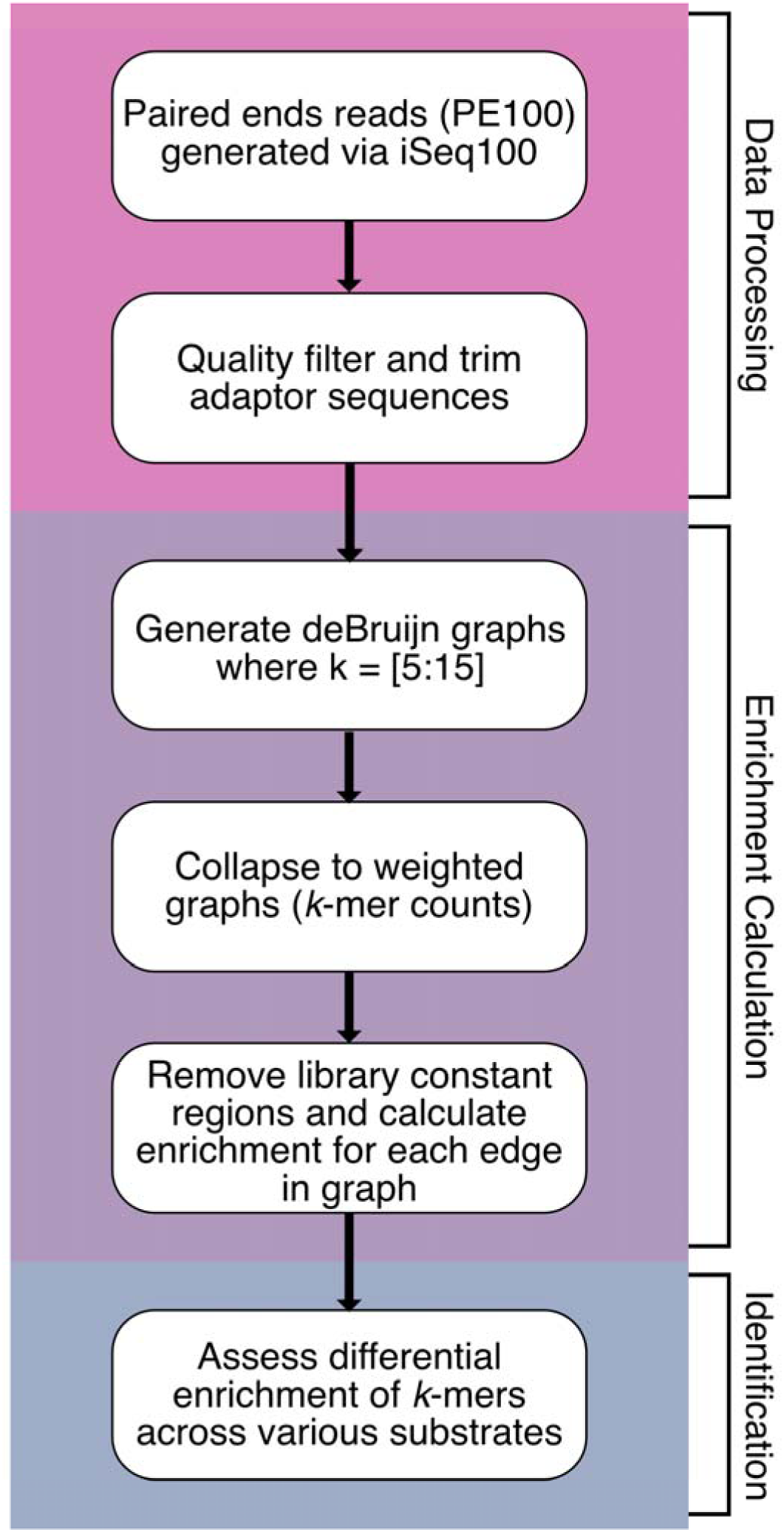
Computational pipeline for reflected nucleic acid *k*-mer space analysis.

**Figure 3.**
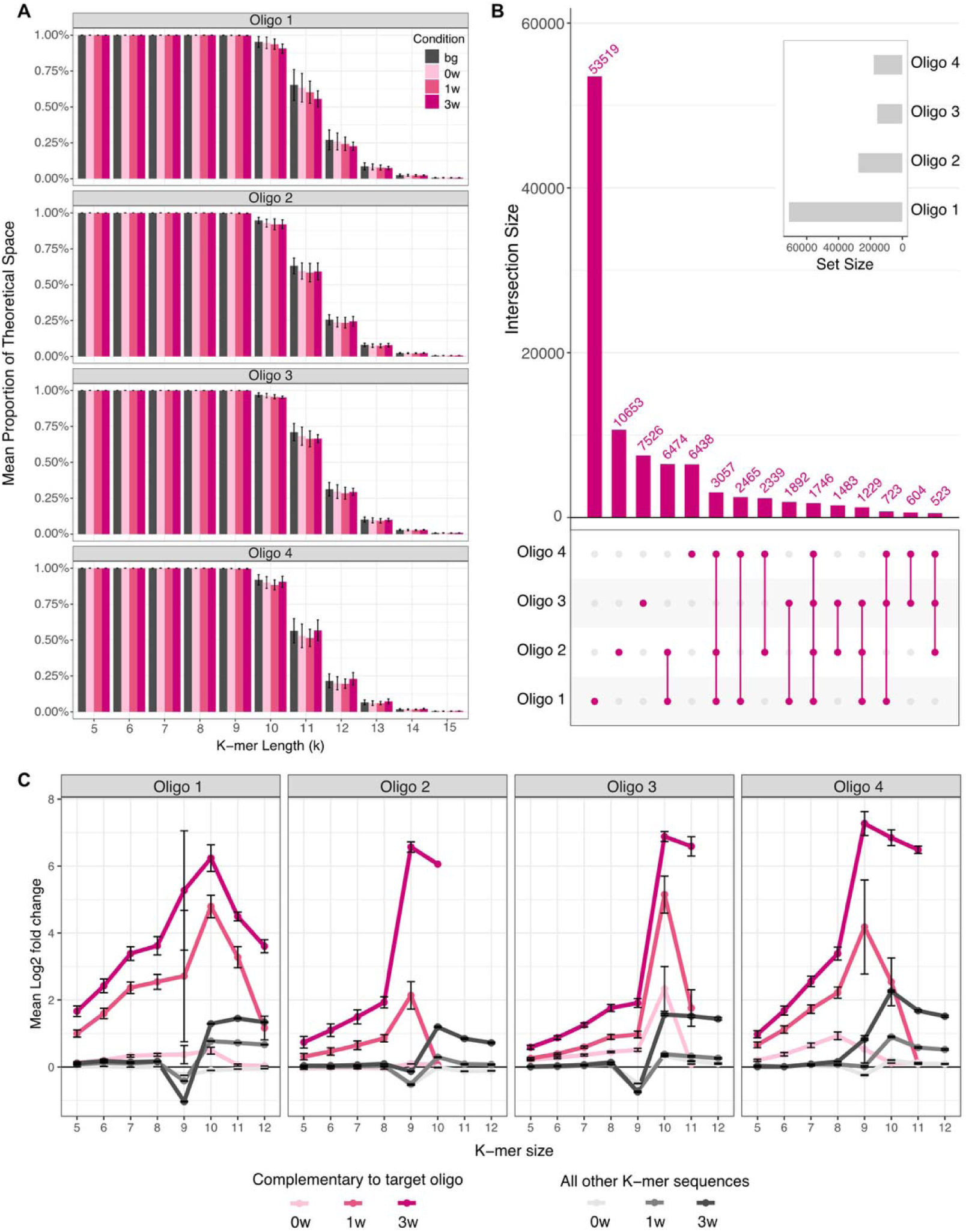
Nature of *k*-mer populations associated with oligonucleotide substrates. (A) Proportion of total theoretical sequence space represented by the measured number of unique *k*-mers for all oligonucleotides, error bars represent standard deviation. (B) Intersection of differentially enriched *k*-mers (5% FDR, *k* = [5:15]) across each of the four oligonucleotide substrates. (C) Effect of washing on representation of enriched *k*-mers at 5% FDR that are fully complementary to the substrate oligonucleotide versus *k*-mers with one or more mismatches with the substrates, where error bars represent standard error.

For *k*-mer sizes 5 through 9, 4*^k^* unique *k*-mers were observed in the library sequences obtained from bg, 0w, 1w, and 3w samples of all four oligonucleotide substrates indicating that 100% of the expected *k*-mer sequence space was represented in the data (**Figure 3A**). For instance, all expected 1024 5-mers and all 2.6 x 10^5^ 9-mers were observed. Since sequencing was capped at 1 million reads and 3 to 7 biological replicates per sample, as the *k*-mer length was increased the proportion of the expected sequence space observed declined progressively. The datasets also demonstrated 90% to 100% coverage of the about 1 million expected 10-mers (**Figure 3A**). For 11-mers the values were between 60% to 95% of possible sequences; for 12-mers 25% to 75%, for 13-mers 12% to 25%, while less than 12% of the expected 0.3 and 1 billion 14- and 15-mers were found, respectively.

Amongst the four oligonucleotide substrates, Oligo 1 data displayed the most coverage at all *k*-mer lengths likely due to the greater number of biological replicates and therefore larger size of the cumulative dataset. With increasing numbers of washes, no reductions were measured in the observed proportions of expected 5- to 9-mers sequence space, while representation of the expected 10- to 15-mer sequence space declined in many 0w, 1w, and 3w samples relative to the bg samples (representative of the starting libraries). However, there were no consistent trends between or within the four oligonucleotide substrates. Taken together, these results indicate a lack of significant skewing of the starting library by synthesis, amplification, or sequencing.

Having validated the diversity of the lightly selected *k*-mer sequence spaces, we hypothesized that washing away of unbound library members should lead to a substantive enrichment of complementary *k*-mers in the population, as opposed to non-complementary *k*-mers. To test this hypothesis the intersections between the sequence space imprints of the four different oligonucleotide substrates were evaluated. To reduce noise, the datasets were first filtered such that only *k*-mers that were found at least 20 times in a dataset were retained.

While several differentially enriched *k*-mers (*k*=[5:15], FDR = 5%) were associated with more than one type of oligonucleotide substrate, significant numbers of *k*-mers were found to uniquely assort with specific oligonucleotide substrate type (**Figure 3B**). The appearance of these uniquely enriched *k*-mers confirms that washing enriches substrate-specific *k*-mer sequences. Indeed, with increasing numbers of washes, complementary *k*-mers consistently demonstrated a corresponding Log2 fold increase in their mean counts relative to their abundance in the starting library (bg) (**Figure 3C**). This was true across all *k*-mer lengths examined but was especially evident at lengths ≥ 9 nt. Overall, these results indicate that despite minimal selection and the presence of background, oligonucleotide specific patterns could be readily distinguished in the sequence space.

While it is perhaps not surprising that specific *k*-mers would be found to associate with oligonucleotide targets, the important point is that the methodology is generalizable. Having developed and validated our experimental and analytical pipeline, we sought to assess the robustness of the pipeline for the classification of non-oligonucleotide materials, including the protein Streptavidin, the polysaccharide Chitin, and the inorganic compound borosilicate glass. Each of these substrates was similarly probed with identical N_15_ ssDNA libraries in quadruplicate and then analyzed using the experimental and computation pipeline described above. The sequencing datasets associated with non-nucleic acid substrates demonstrated similar coverage of the expected *k*-mer space as was observed with the oligonucleotide substrate-bound libraries (**Supplementary Figure 2**).

Similar to the association of unique *k*-mer subsets with the oligonucleotide substrates, borosilicate glass, Chitin, and Streptavidin were also found to associate with unique sets of non-overlapping *k*-mers (**Figure 4A-C**). To assess whether these broad differences in *k*-mer substrate associations were indeed reflective of distinctive substrate-specific *k*-mer enrichment, positively enriched *k*-mers in the 0w, 1w, and 3w sequence imprints selected by Streptavidin, Chitin, borosilicate glass, and the original four oligonucleotide substrates were subjected to non-metric multidimensional scaling (NMDS) analysis. NMDS ordination based on Bray-Curtis dissimilarity demonstrated excellent fit (stress = 0.087; non-metric R^2^ = 0.992) and showed distinct substrate-specific clusters (**Figure 4C**). Analysis of similarities (ANOSIM) indicated statistically significant separation between the seven substrate-associated clusters (R = 0.981 and p = 0.001; **Supplementary Figure 3**). Permutational multivariate analysis of variance (PERMANOVA) verified that substrate type could explain 78.6% of the total variation (F = 8.58 and p = 0.001; **Supplementary Figure 3**) while analysis of multivariate homogeneity of groups dispersions (betadisper) confirmed that variances within the different substrate groups were not statistically different (F = 0.279 and p = 0.937; **Supplementary Figure 3**).

**Figure 4.**
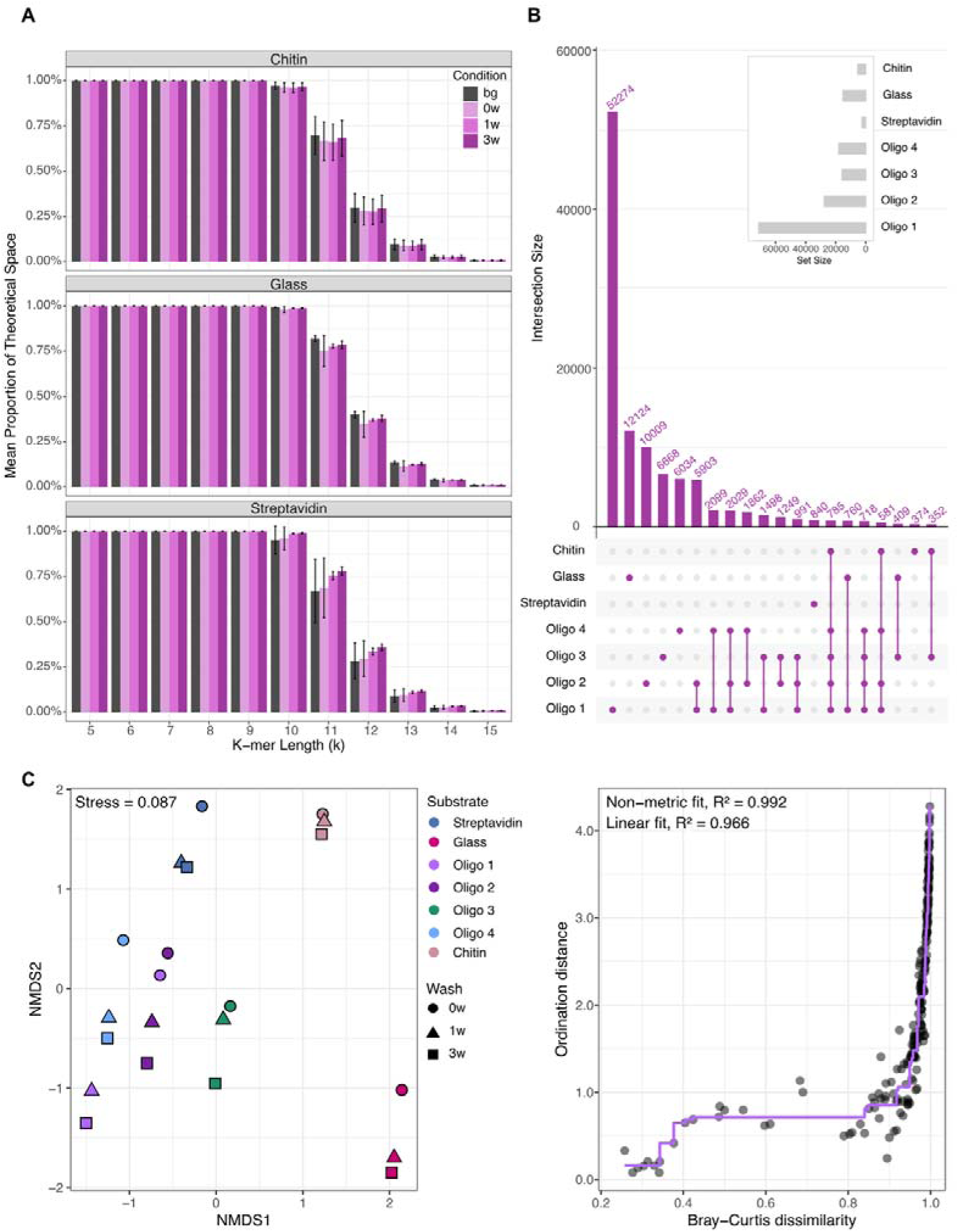
Nonmetric multi-dimensional scaling analysis of substrates. (A) Proportion of total theoretical sequence space represented by the measured number of unique *k*-mers for Chitin, Glass, and Streptavidin, error bars represent standard deviation. (B) Intersection of differentially enriched *k*-mers (5% FDR, *k* = [5:15]) across each substrate. (C) NMDS clustering of enriched *k*-mers across substrates. Ordination fit of NMDS is shown in the right panel.

To further verify the specificity of substrate-associated *k*-mer spaces we posited that de Bruijn graphs of the enriched *k*-mers would map uniquely to the corresponding substrates. To test this hypothesis, the top 100 most enriched *k*-mers of sizes 5-15 nt in the 3w datasets were used to build contiguous sequences. For each substrate, contiguous sequences exhibiting the longest median length per *k*-mer size per substrate (**Supplementary Figure 4**) were combined and analyzed using hierarchical clustering and BLAST. Ultimately, this resulted in analyzing contiguous sequences derived from 8-mers for all substrates except Oligo 4, Streptavidin, and Chitin which used 7-mers, 9-mers, and 14-mers, respectively. Contigs were hierarchically clustered using *k*-mer cosine distance metrics (*k*=5) and were observed to cluster together based on the substrate of origin, with some overlap between Chitin and Streptavidin contigs (**Figure 5**). Overall, contigs for individual substrates were more closely related to each other than to sequences from another type of substrate. This demonstrates that low affinity binding species can be used to agnostically distinguish (‘decode’) different molecular surfaces that they come in contact with.

**Figure 5.**
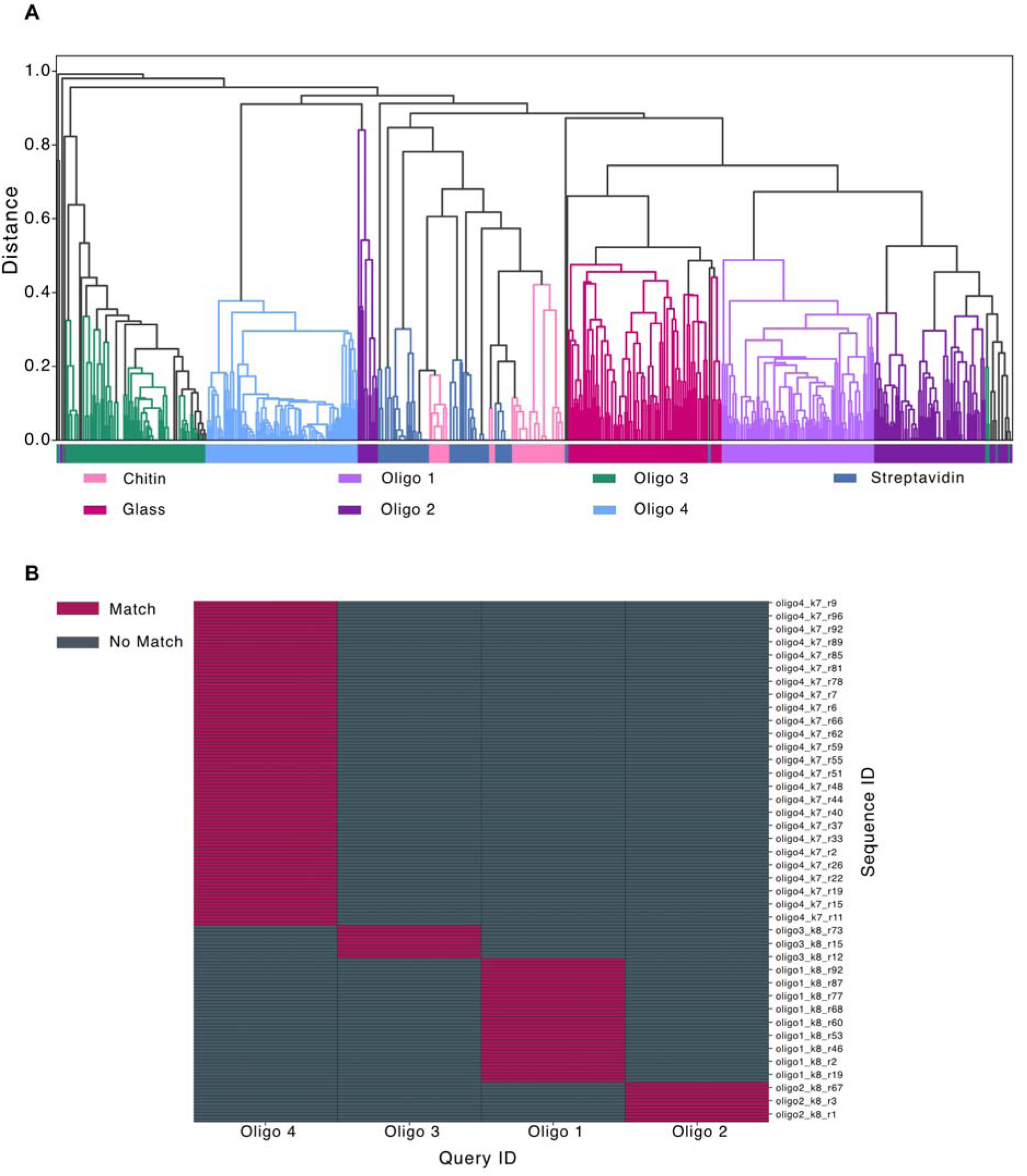
Contig sequence clustering reflects substrate identity. (A) *K*-mer cosine clustering of contiguous sequences built from top 100 enriched *k*-mers. Results are visualized as a dendrogram and colored by substrate name. (B) Oligonucleotide substrates aligned to contig sequences using BLAST, and overall patterns visualized as a heat map.

Substrate specificity was then evaluated by performing BLASTn analysis using the oligonucleotide substrate sequences as queries against data sets containing de Bruijn graph-generated contig sequences obtained from all seven datasets. Satisfyingly, the oligonucleotide substrates were found to align only with their own enriched contig sequences (**Figure 5**). In contrast, contig sequences generated from *k*-mers enriched by borosilicate glass, Streptavidin, or Chitin did not align with any of the four oligonucleotide substrates. Conversely, when reference-free consensus sequences were built from the de Bruijn contigs associated with each of the seven substrates (**Supplementary Table 2**), 48% to 100% of the four original oligonucleotide substrate sequences were specifically reconstructed only by their respective consensus sequence (**Supplementary Table 3**). These results fully demonstrate that analysis of *k*-mer space for weakly associated nucleic acid species can not only distinguish oligonucleotide substrates from non-nucleic acid substrates but can even capture distinctions between otherwise closely related chemicals (oligonucleotides).

It is tempting to speculate that decoding surfaces as DNA imprints may reveal differences in surface physicochemistry. While borosilicate glass enriched a diversity of *k*-mers, there was a predominance of sequences containing T, perhaps because thymine is one of the most hydrophobic nucleobases and therefore might provide increased opportunities for hydrophobic interactions.^16,17^ Similarly, cytosine was also preferred in sequences binding glass, potentially because this nucleotide could support electrostatic and hydrogen bonding interactions with glass.^16,18^ Finally, the *k*-mers enriched by chitin or streptavidin tended to be longer and have higher frequencies of A, potentially driven by its ability to form CH-Pi stacks with carbohydrates.^19–22^ Overall, though, the true value of our method is that none of these explanations matter or are necessary: the sequences themselves provide an agnostic readout of the surfaces they contact.

## CONCLUSIONS

Given the specificity and strength of DNA hybridization as well as the vast prior art on discovery of aptamers that can bind a variety of non-nucleic acid substrates, we hypothesized that substrate-specific ssDNA library members would be somewhat enriched even in lightly selected samples. However, whether statistically significant enrichment would be readable against a highly noisy background, due to the minimal selection, to allow substrate-level resolution of DNA binding patterns was an open question. Our results now demonstrate that substrate-specific de Bruijn *k*-mer imprints can be captured and detected in noisy DNA binding data not only for oligonucleotide substrates but also for a wider chemical and structural variety of substrates whose interaction with DNA libraries is independent of direct nucleobase pairing.

One limitation of the current experimental approach is the use of particulate substrates and amplification of their bound library members by direct PCR analysis where the whole substrate:DNA complex is added to the PCR reaction. However, sample preparation methods may be readily adjusted using existing tools to enable analysis of a wider and varied range of substrates. For instance, to analyze particulate materials that might inhibit PCR, substrate-selected library members can be eluted from the substrates by heat denaturation or by using chaotropes followed by column-based cleanup of the recovered DNA prior to PCR amplification. Similarly, library members bound to non-particulate substrates that cannot be readily collected by centrifugation could be separated from unbound DNA by various methods, such as size exclusion chromatography, surface immobilization of the substrates, or filtration, prior to PCR analysis. In fact, this ability to select, recover, and amplify substrate-bound DNA should allow analysis of even very dilute samples, which may elude evaluation by other methods. Ultimately, our deliberate minimization of experimental steps and use of high-throughput pipelines should facilitate process mechanization for remote and automated implementation.

Analysis of the data revealed imprints specific to each substrate. De Bruijn graph-based *k*-mer analysis has been used extensively for analysis of complex genetic data and discovery of nucleic acid biomarkers, such as in identifying group-specific sequences in metagenomic data, creating reference-free representations of genomes and predicting antibiotic resistance phenotypes, and identifying non-reference sequence variants in genomes.^23–25^ We now demonstrate for the first time that the information contained in the unique enrichment patterns of the de Bruijn graph-based DNA *k*-mer imprint of a material can serve as an independent measure to distinguish materials, including those of non-nucleic acid composition. The positively enriched *k*-mer imprints of the seven different tested substrates were significantly different (ANOSIM: R = 0.981 and p = 0.001) with substrate type explaining 78.6% of the total variation. Furthermore, imprints derived from nucleic acid substrates could agnostically reconstruct 48% to 100% of their original substrate.

Our results now set the stage for applying de Bruijn *k*-mer analysis for high-throughput, agnostic material analysis and classification using the readily derived unique nucleic acid sequence space imprints. This method and computational algorithm should be generalizable to the analysis of materials with other chemistries, including a variety of polymer materials. Indeed, the ability to define sequence (ala oligonucleotide analyses) may carry over to the more precise identification of specific compositions (e.g., block co-polymers) or even sequences. Conversely, sequence-based libraries beyond oligonucleotides, such peptide or polyurethane libraries,^26^ could potentially themselves be used to deconvolute materials.

## METHODS

### Chemicals and reagents

All chemicals were purchased from Sigma-Aldrich (St. Louis, MO, USA) unless otherwise noted. All molecular biology enzymes and magnetic beads were obtained from New England Biolabs (Ipswich, MA, USA) while NextGen sequencing cartridges and reagents were acquired from Illumina (San Diego, CA, USA). Ahlstrom Grade 161 binder-free microglass fiber filter paper (1610-0210) was purchased from VWR (Radnor, PA, USA). Oligonucleotides were synthesized by Integrated DNA technologies (Corralville, IA, USA). The N15 randomization in the DNA library was performed by hand-mixing the four standard deoxyribonucleotide phosphoramidites in equimolar ratio. The resulting library was purified following separation by polyacrylamide gel electrophoresis.

### Material DNA imprinting protocol

*Substrate preparation:* The hemiduplex oligonucleotide substrates 1, 2, 3, and 4 (**Supplementary Table 1**) were prepared by annealing 10 µM of biotinylated Oligo 1, 2, 3, and 4, respectively, with 10 µM of Oligo dT in 1X PBSM (137 mM NaCl, 2.7 mM KCl, 10 mM Na_2_HPO_4_, 1.8 mM KH_2_PO_4_, 5mM MgCl_2_, pH 7.4) by incubation at 95 °C for 1 min followed by slow cooling (0.1 °C/sec) to room temperature. One microliter (4 µg) of streptavidin-coated magnetic beads (NEB; catalog number S1420S) were washed twice with 20 µL of 1X PBSM and then incubated with 10 µL of either 1X PBSM buffer alone or 1X PBSM containing 100 picomoles of a biotinylated hemiduplex oligonucleotide substrate. The bead suspensions were placed on a rotator for 30 min at room temperature to enable immobilization of the biotinylated hemiduplex DNA. Subsequently, beads were collected using a magnetic rack (NEB; catalog number S1509S) and the supernatants were discarded. Beads were washed five times with 20 µL of 1X PBSM prior to application of the LibProbe12 N_15_ ssDNA library. Glass fiber filter paper discs (3 mm diameter) and 5 µL of chitin-coated magnetic beads (NEB; catalog number E8036S) were prepared for LibProbe12 DNA library binding by washing thrice with 20 µL of 1X PBSM.

Oligonucleotide substrate annealing reactions were performed in 0.2 mL thin-walled PCR tubes (Axygen-Corning, Corning, NY, USA). All other manipulations were performed in 1.5 mL or 2 mL DNA LoBind microtubes (Eppendorf, Hamburg, Germany).

### Single stranded DNA N_15_ probe library preparation and substrate binding

A 10 µM solution of the single stranded DNA library (LibProbe12) with a N_15_ randomized region (N_15_ ssDNA) in 1X PBSM buffer (137 mM NaCl, 2.7 mM KCl, 10 mM Na_2_HPO_4_, 1.8 mM KH_2_PO_4_, 5mM MgCl_2_, pH 7.4) was subjected to refolding by incubation at 95 °C for 1 min followed by slow cooling (0.1 °C/sec) to room temperature. Two picomoles each of the refolded N_15_ ssDNA library were applied to four identical preparations of each substrate per experimental replicate. Library application was performed in either 10 µL (beads) or 3 µL (glass paper discs) volume of 1X PBSM. The substrates were incubated with the LibProbe12 library for 30 min on a rotator at room temperature. Subsequently, these quadruplicate DNA library-treated substrates in each experiment were prepared for PCR amplification of the DNA library members in one of the following four ways: (i) One substrate:LibProbe12 library mixture, designated as ‘bg’, was directly added to the PCR reaction without further processing. These samples retained all the LibProbe12 library members that were initially applied to the substrate. (ii) For the second substrate:LibProbe12 library mixture, designated as ‘0w’, the supernatant containing unbound LibProbe12 library members was removed after collecting the magnetic beads or the glass paper discs using a magnetic rack or brief centrifugation, respectively. The retained glass paper disc was directly added to the PCR reaction while the reserved magnetic beads were resuspended in 10 µL of 1X PBSM before being transferred into the PCR reaction. (iii) The third substrate:LibProbe12 mixture, designated as ‘1w’, underwent similar removal of unbound library members as performed for the ‘0w’ samples. In addition, these substrates were washed once with 20 µL of 1X PBSM prior to transfer into PCR tubes as described above. (iv) The fourth substrate:LibProbe12 mixture, designated as ‘3w’, underwent similar removal of unbound library members as performed for the ‘0w’ samples. In addition, these substrates were washed thrice with 20 µL of 1X PBSM prior to transfer into PCR tubes as described above. Library refolding was performed in 0.2 mL thin-walled PCR tubes (Axygen-Corning) while library incubation with substrates was performed in 1.5 mL or 2 mL DNA LoBind microtubes (Eppendorf).

### PCR amplification and Illumina sequencing

LibProbe12 library members in the bg, 0w, 1w, and 3w samples were both amplified and appended with Illumina sequencing adaptors in a single 100 µL one-pot nested PCR reaction containing 1X ThermoPol buffer (NEB; catalog number B9004S; 20 mM Tris-HCl, 10 mM (NH_4_)_2_SO_4_, 10 mM KCl, 2 mM MgSO_4_, 0.1% Triton® X-100, pH 8.8) supplemented with 0.2 mM deoxyribonucleotide mix and 5 units of Taq DNA polymerase (NEB). The PCR reactions also received 0.04 µM each of the inner PCR primers NGS.1F and NGS.2R, 0.4 µM of the i5 indexed outer forward primer NGSTSBC01F, and 0.4 µM of either NGSTSBC05R (bg samples), NGSTSBC06R (0w samples), NGSTSBC07R (1w samples), or NGSTSBC08R (3w samples) i7 indexed outer reverse primers (**Supplementary Table 1**). The PCR reactions were held at 95 °C for 2 min followed by 30 cycles of 10 seconds incubation at 95 °C, 15 seconds incubation at 55 °C, and 30 seconds incubation at 72 °C. The PCR amplicons were gel purified, quantitated, and diluted in 10 mM Tris pH 8.5 prior to preparing a pooled library containing 40 pM each of the bg, 0w, 1w, and 3w amplicons from a substrate along with 40 pM of PhiX v3 control DNA (Illumina). The pooled libraries were sequenced on an iSeq-100 platform (Illumina) using paired-end 100 cycle sequencing according to the manufacturer’s instructions.

### Data processing and analysis

#### Sequencing QC and initial processing

Custom Python and R scripts were used unless otherwise indicated. All code can be found on GitHub at https://github.com/rachaelcox/dna_fingerprints.git. All sequencing data were subjected to trimming, filtering, and merging using fastp v0.23.4 (**Figure 2**).^27^ Sequencing data was visualized before and after quality control using MultiQC v1.19 to confirm adaptor removal, low quality read removal, and correct library sequence structure (i.e., constant flanks with an interior degenerate sequence).^28^

#### k-mer generation

We generated de Bruijn graphs from each of the processed data sets for values of *k* ranging from 5 to 15, netting 11 total de Bruijn graphs per replicate. To construct a weighted graph, we counted each unique *k*-mer edge that appeared in each dataset while maintaining directedness of the graph. We compared the number of unique *k*-mers recovered at each value of *k* in *k* = [5:15] to the maximum unique *k*-mers that could be expected at each length (e.g., where *k* = 5 for a de Bruijn graph of sequences comprised of 4 nucleotides, the total number of unique *k*-mers should not exceed 4^5^ = 1,024). After a 20 count cutoff and subtraction of *k*-mers originating from the constant regions of the library members, the weighted graphs were used to compute enrichment of *k*-mers in unwashed (“0w”), once-washed (“1w”), and thrice-washed (“3w”) samples in comparison to counts of those *k*-mers in the corresponding background (“bg”) data. This enrichment was computed using the limma package for differential gene expression as implemented in degust.^29,30^

#### Data visualization

Enriched *k*-mers that are specific to individual substrate or combination of substrates were visualized using the ComplexUpset (v.1.3.3) and UpsetR (v1.4.0) R packages^30–32^ .The effect of washing on enriched *k*-mers was also analyzed by plotting the mean log2 fold change for complimentary and non-complimentary *k*-mers at each length of *k* for each oligonucleotide. These results are visualized as a line plot in R where error bars correspond to the standard error of the mean.To assess whether the enriched pools were sufficiently diverse to differentiate substrates agnostically, custom R scripts were used to extract the most enriched *k*-mers at a 5% FDR for each substrate which was then used as input for non-metric multidimensional scaling (NMDS) analyses.^31^ Dissimilarity between enriched *k-*mers was calculated using enriched 10-mer data. The Bray Curtis metric was used over Euclidean estimations to maintain the rank order of *k-*mers. Ordinations and stress plots were visualized using the vegan package (v.2.7.1) in R. ANOSIM, PERMANOVA, and betadisper analyses of the clustering were performed using the Vegan package with the number of permutations set to 999. The top 100 most enriched 5- to 15-mers in the 3w datasets were also used to assemble contiguous sequences. The median sequence length of the reconstructed sequences of each *k*-mer size across all substrates were visualized using box plots.

#### Clustering analysis

Contiguous sequences associated with the highest median sequence length were extracted and concatenated into a single FASTA file. The resulting sequences were hierarchically clustered using *k-*mer cosine distances via the scikit-learn library (v1.7.2), using *k* = 5. Clusters were visualized as a dendrogram and colored by substrate name using the SciPy (v1.15.2) and MatplotLib (v3.10.6) libraries. As expected, the clustering supports target-specific classification of the various *k*-mer-built contiguous sequences.

#### BLAST analysis

In addition, the contiguous sequences were used to perform BLAST analyses against the oligonucleotide substrates. The BLAST analysis was performed by creating a custom Query database containing the four oligonucleotide substrate sequences Oligo 1, 2, 3, and 4.^32^ Blast was performed using blastn (v.2.12.0) with default parameters (reward 2; penalty −3; gapopen 5; gapextend 2; evalue 0.05; dust yes; and soft_masking true) with word_size 7. The resulting alignments were visualized as a heatmap using seaborn (v0.13.2).

#### Nucleotide frequency of k-mers

Nucleotide frequency of *k*-mer sequences was determined by processing data extracted from the *k*-mer weights file generated for each substrate at each *k*-mer length. Analysis was performed on sequences in which the length of the sequences was greater than one. Individual nucleotide frequencies were calculated by summing the total instances of individual base pairs (A,C,T,G) in each sequence and divided by the total sequence length. Data was processed using the data.table (v.1.17.8) and stringer (v.1.5.1) packages. Results were visualized using the grid (4.3.3), and ggplot2 (v.4.0.0) packages.

#### Reference-free consensus assembly

Geneious Prime® 2023.0.4 ‘Multiple Align’ tool was used to generate reference-free consensus sequences from the contiguous sequences associated with the highest median sequence length. The contiguous sequences were aligned using in-built Geneious global alignment with free end gaps, 93% similarity cost matrix, default gap open and extension penalties of 12 and 3, respectively, and 5 refinement iterations. Specific reconstruction of original bait oligonucleotides was verified using BLASTn with default settings adjusted for word size of 7.

## Acknowledgements

A.D.E. acknowledges funding from the National Aeronautics and Space Administration (c) and the Welch Foundation (F-1654). S.B. acknowledges the Texas Proof-of-Concept award from the University of Texas at Austin Discovery to Impact. E.M.M. acknowledges funding from the National Institute of General Medical Sciences (R35GM122480) and the Welch Foundation (F-1515). The authors acknowledge additional support generously provided by Tito’s Handmade Vodka. Computational analyses were performed using the Biomedical Research Computing Facility at UT Austin, Center for Biomedical Research Support (RRID#: SCR_021979). The authors would also like to thank Advanced Micro Devices for the donation of critical hardware resources from its HPC fund.

