## Supplementary files for "Agnostic material classification using differential de Bruijn graphs of DNA imprints"

Supplementary Table 1

Supplementary Figure 1

Supplementary Figure 2

Supplementary Figure 3

Supplementary Table 2

Supplementary Table 3

Supplementary Figure 4

**Supplementary Table 1. Sequences of oligonucleotides used in this study.**

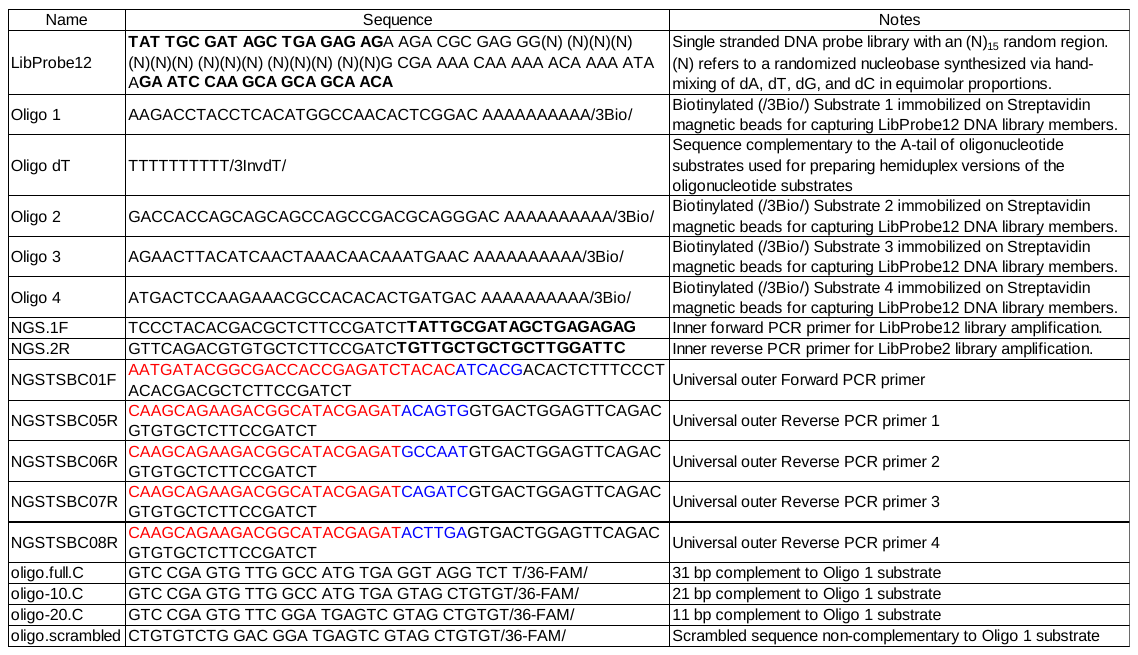

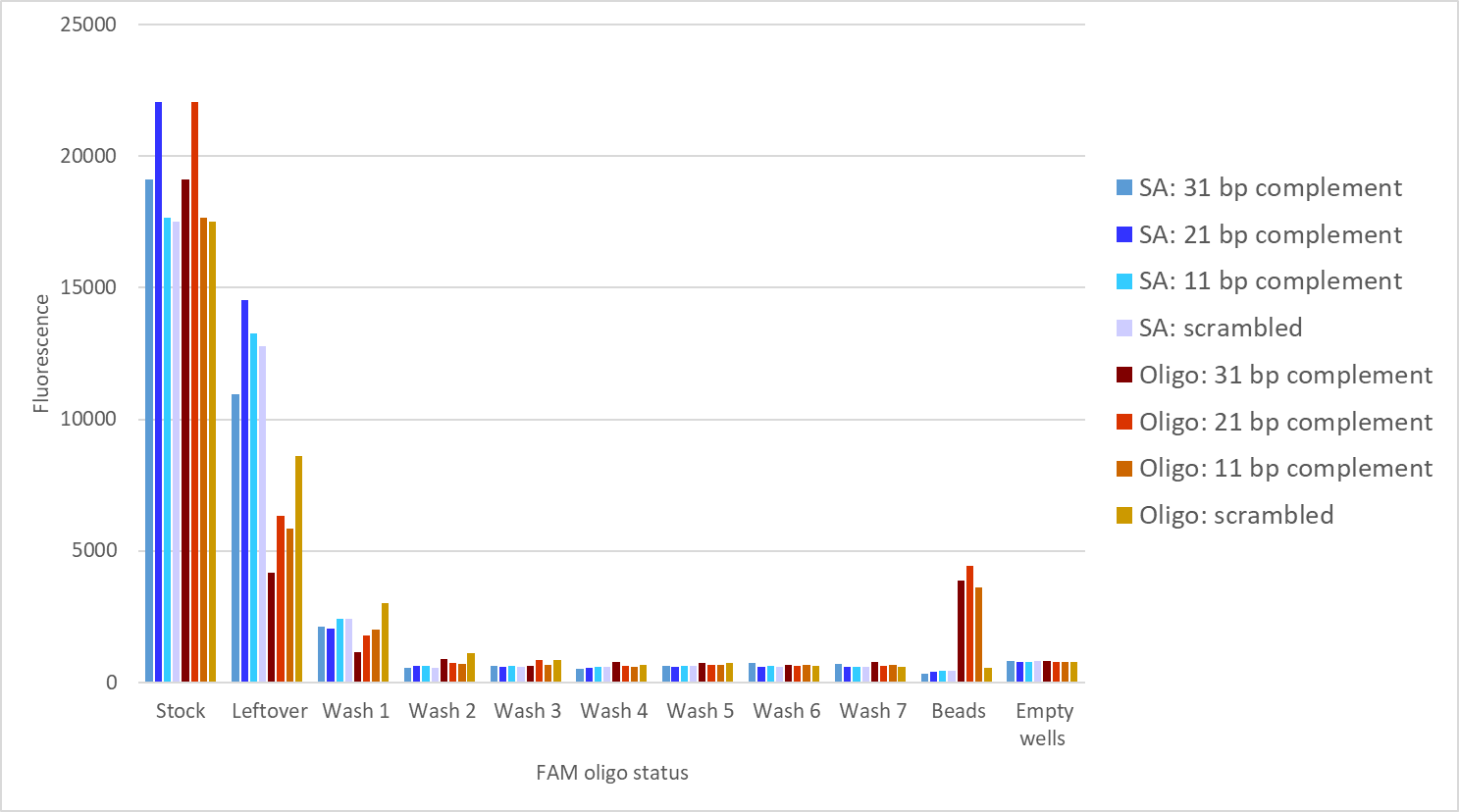

**Supplementary Figure 1. Impact of washing on retention of substrate-bound oligonucleotides.** Streptavidin-coated magnetic beads (‘SA’) displaying immobilized hemiduplex oligo 1 DNA substrates (‘Oligo’) were incubated with two picomoles of non-complementary (‘scrambled’), fully complementary (‘31 bp complement’) or partially complementary (‘21 bp complement’ and ‘11 bp complement’) fluorophore-labeled probe oligonucleotides. Endpoint measurements of fluorescence in probe samples taken before application to beads (‘stock’), leftover after bead binding (‘leftover’), removed in wash buffer after indicated number of washes (‘Wash 1 to Wash 7’), retained on beads (‘beads’) are depicted. Background noise measured in empty wells of the same microplate are reported as fluorescence in ‘empty wells’.

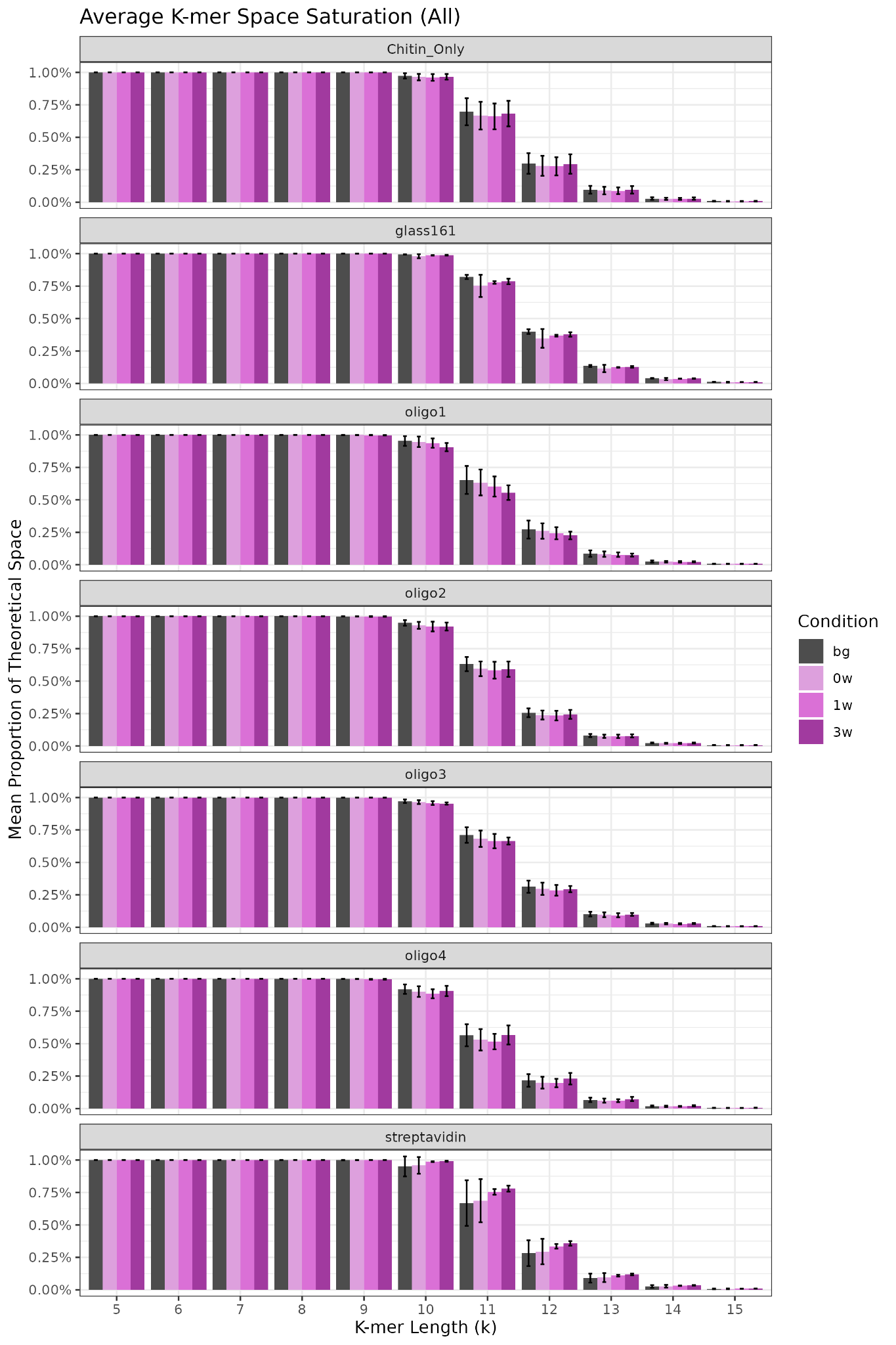

**Supplementary Figure 2. Average expected *k*-mer space representation and retention during analysis of non-nucleic acid and nucleic acid substrates.**

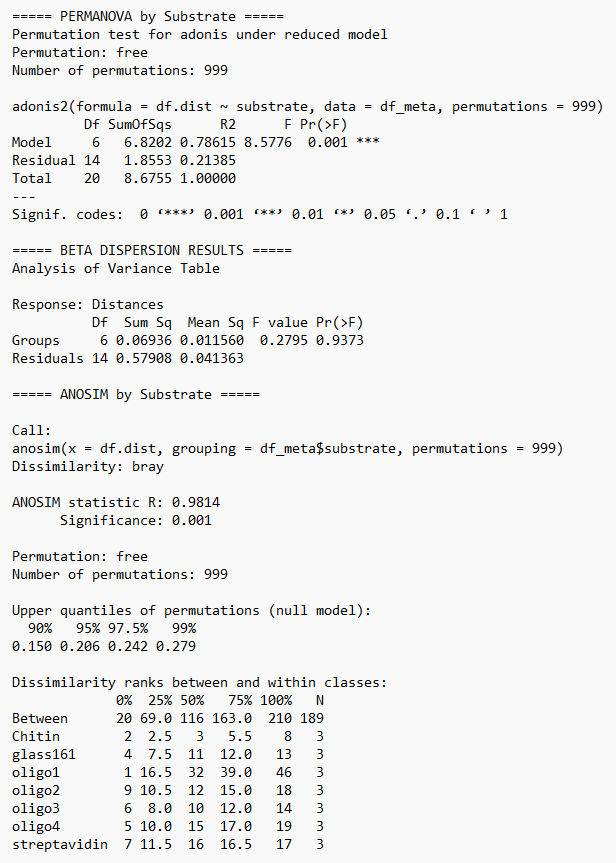

**Supplementary Figure 3. Statistical analysis of substrate-specific clustering of positively enriched *k*-mers.**

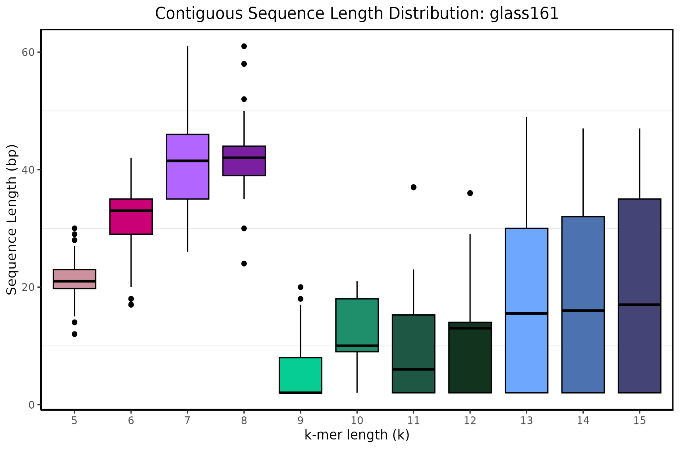

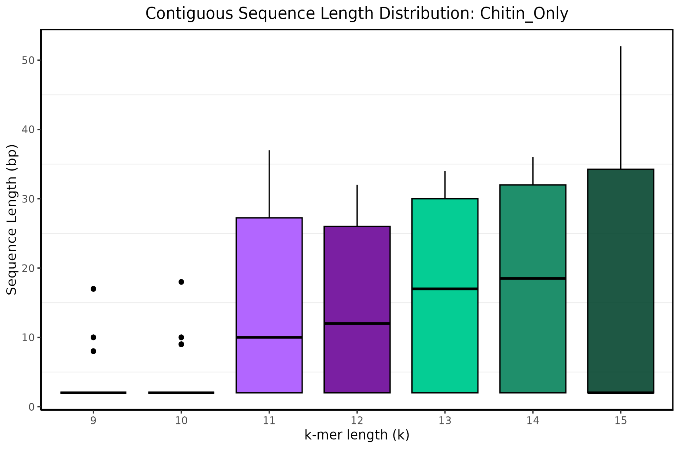

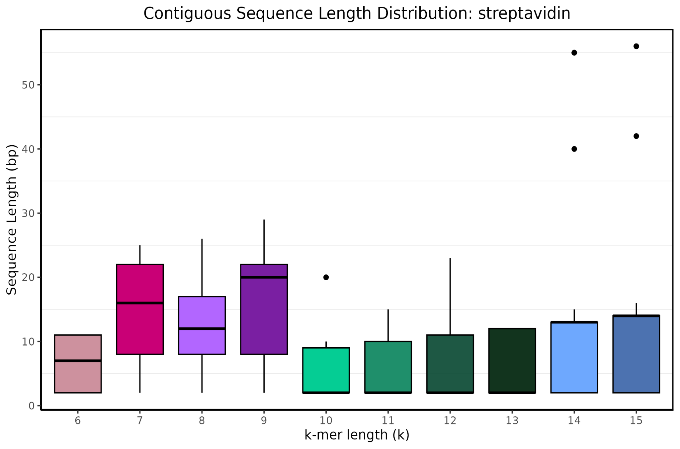

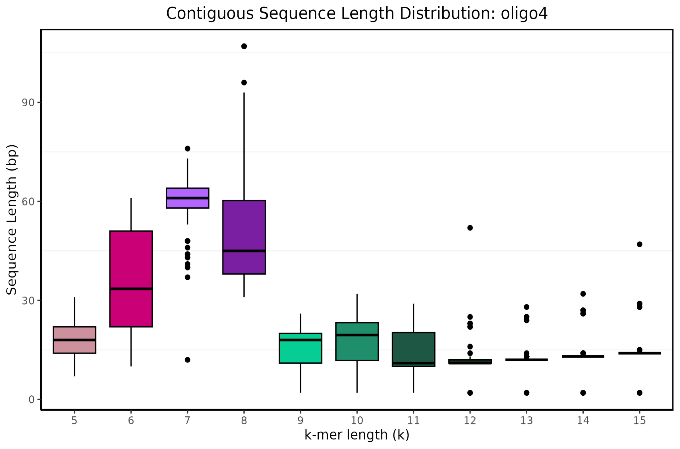

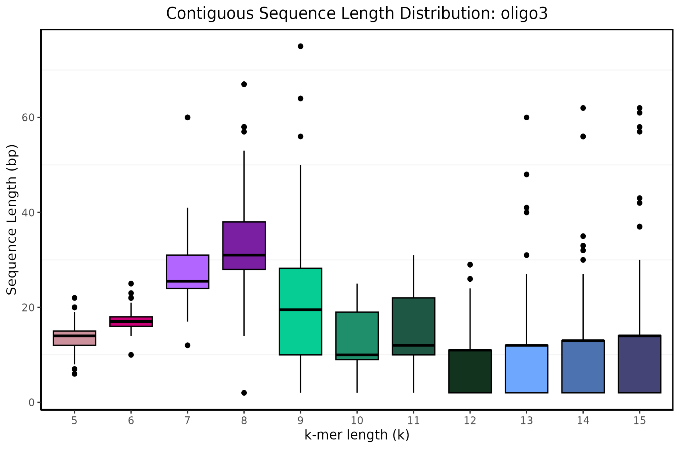

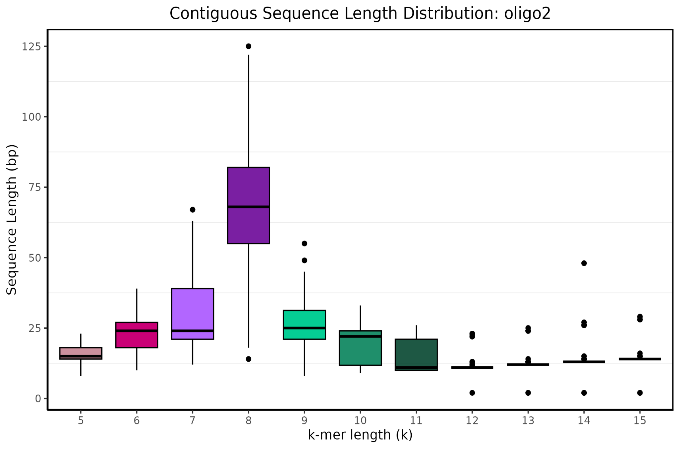

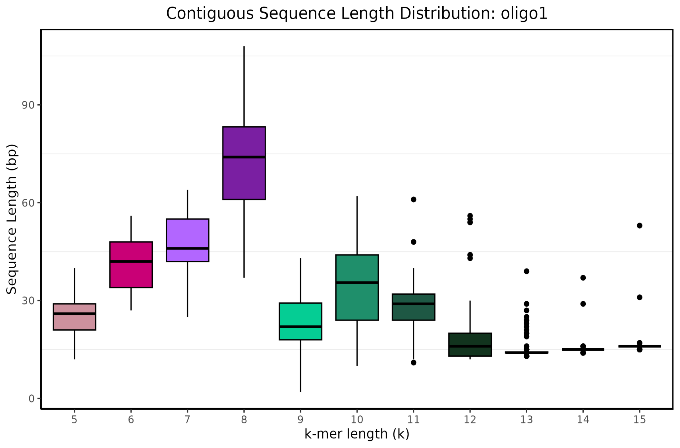

**Supplementary Figure 4. Median contiguous sequence length for each *k*-mer size.**

**Supplementary Table 2. Reference-free consensus of substrate-associate contiguous sequences.**

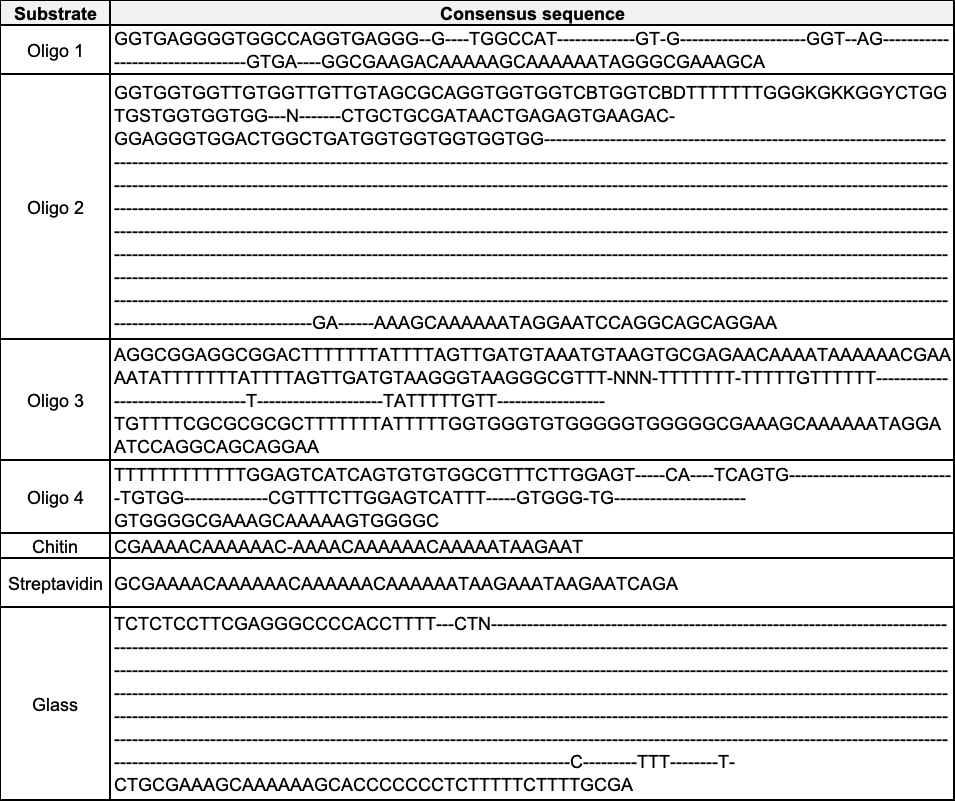

**Supplementary Table 3. Identification of oligonucleotide substrate sequence information captured by the reference-free consensus sequences using BLAST analysis.**

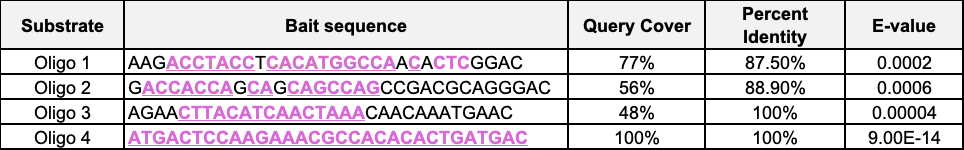

Captured oligonucleotide substrates sequences are denoted in pink. Query Cover: percent of the substrate sequence recaptured by the consensus.
